# Kin selection for cooperation in the honey bee microbiome

**DOI:** 10.64898/2026.11.12.766218

**Authors:** Laurence J. Belcher, Rosie Randall, Sarah G. Flint, Harvey J. Jeffrey, Zohar Katz, Anna E. Dewar, Stuart A. West

**Affiliations:** Department of Biology, University of Oxford, Oxford, OX1 3SZ, United Kingdom; School of Biosciences, The University of Sheffield, Western Bank, Sheffield, S10 2TN, United Kingdom; St John’s College, Oxford, OX1 3JP, United Kingdom

**Keywords:** kin selection, cooperation, microbiome, honey bee, social evolution, molecular evolution

## Abstract

Bacteria engage in many cooperative behaviours which provide benefits to other cells in the population. Lab experiments have suggested that these costly behaviours can be favoured by kin selection due to the benefits they provide to closely related cells. However, we know much less about the role of kin selection in natural populations. Here, we used two complementary methods to measure and compare kin selected signatures of cooperation in 19 species of bacteria found in the guts of honey bees. We used bioinformatic tools to count the number (and proportion) of genes for cooperative traits in a genome, and used molecular population genetics to measure how natural selection is acting on those genes. We found evidence for a consistent signature of kin selection for cooperation on genes controlling cooperative traits across species. Genes for cooperative traits had more non-synonymous relative to synonymous polymorphism than genes for private traits, had more protein-changing mutations and more codons under positive selection, and were lost more often. From this signature, we estimate an average relatedness of 0.73 between interacting cells. However, we found no evidence supporting the prediction that this signature varies with relatedness estimated from metagenomes. These patterns suggest that whilst genes for cooperative traits are undergoing kin selection in natural populations, there may be other factors influencing how the action of kin selection varies across species.

**Impact Statement:** Bacteria cooperate by secreting molecules that benefit other cells, such as enzymes that break down food and molecules that take up iron. Kin selection theory predicts that genes for these cooperative traits are under weaker selection than genes for private traits, which benefit only the cell that makes them. We tested this prediction across 19 species of bacteria from the honey bee gut. As predicted, we found that genes for cooperative traits had more non-synonymous relative to synonymous polymorphism than those for private traits. Between strains of the same species, genes for cooperative traits had more protein-changing mutations and more positively selected codons, and were lost more often. However, these differences did not vary across species with relatedness estimated from metagenomes. One possible reason is that bacteria interact in small patches of the gut, so our whole-gut metagenomic estimates of relatedness may not be accurate.

## Introduction

Bacteria live social lives [1–4]. They cooperate with each other to secrete a remarkable array of molecules allowing them to engage in foraging for nutrients, competitive rivalries, and the neutralisation of antibiotics [5–8]. These cooperative ‘public goods’ are costly to produce but can be favoured by kin selection if they provide benefits to relatives who share the gene for cooperation [9]. Interactions between relatives are likely to be common in bacteria because secreted molecules mostly provide benefits to nearby cells, who are likely to be relatives due to clonal growth [10].

Laboratory experiments have suggested that kin selection can explain cooperation in bacteria [5,10]. However, we know much less about bacterial cooperation in natural populations, and it is unclear whether lab conditions are representative of natural environments [4,11]. It can also be difficult to know what the natural environment of a species is, and we cannot always assume that where we find a species is where it lives. For example, almost 400 species of bacteria can be found on $1 bills [12]. A few species have been well-studied in their natural environment, such as the opportunistic pathogen *Pseudomonas aeruginosa* which can cause chronic infections in humans [13,14], and *Pseudomonas fluorescens* and related species in soil and freshwater [15,16]. However, we lack sufficient knowledge from enough species to test theory across species.

One key prediction from kin selection theory is that as relatedness increases, we should see more cooperation. This prediction has been widely supported in animals, both within species (kin discrimination) and with across species comparative studies [17,18]. In contrast, this prediction has rarely been tested in natural populations of bacteria or other microorganisms. In an innovative study, Simonet & McNally [19] provided a test across 101 bacterial species that form part of the human microbiome. They compared genomic similarity within and between hosts to estimate relatedness for each species and used bioinformatics tools to count the number of genes for cooperative traits in each species’ genome. They found that species with higher relatedness had more genes for cooperative traits.

The generality of this result is unclear. Is having more genes for cooperative traits a good measure of how much cooperation is occurring at the species level? Is the number of genes the key measure, or is it the percentage of genes that are for cooperative traits? Could we instead predict that the total number of genes for cooperative traits stays the same, but kin selection on them gets stronger? In addition, previous methods for identifying genes for cooperative traits may have missed many of these genes [20]. Could this have biased the results?

An alternative approach is to look for a genomic signature of kin selection in genes for cooperative traits [21–26]. If a gene is evolving under direct selection, it gives a direct (personal) benefit to an individual expressing the gene. If a gene is evolving under kin selection, it gives an indirect benefit to another individual. This individual has a certain likelihood of sharing the gene for cooperation, determined by relatedness. When relatedness is less than one, the gene sometimes provides benefits to individuals that do not share the gene, which causes selection on the gene to be relaxed. The degree to which selection is relaxed is proportional to relatedness, as this determines how often the gene gives benefits to individuals that do not share the gene. Population genetic theory predicts that when selection is relaxed, deleterious mutations are less likely to be removed by purifying selection, and positive selection is less effective, so beneficial mutations are less likely to fix. Consequently, when relatedness is less than one, genes for cooperative traits that are favoured by kin selection are predicted to have both increased nucleotide polymorphism and increased sequence divergence, relative to genes for private traits, which evolve under direct selection (Figure S1). We have previously applied this method in two species of bacteria, finding a signature of kin selection for cooperation in genes that were thought to be for cooperative traits from laboratory studies [14,27]. However, this method has never been applied to compare selection across species.

A useful system to test the prediction that higher relatedness leads to more cooperation is provided by the microbiome of the honey bee *Apis mellifera*. Due to the honey bee’s huge economic importance for pollination and a rapid deterioration in colony health, there is large interest in understanding and engineering the microbiome. Recent metagenome sequencing from multiple worker bees in a single colony provides a natural population to study how selection is acting on genes for cooperative traits across bacterial species [28]. At the onset of adulthood, workers acquire their gut microbiome through social transmission from other workers [29]. Each bee carries only a subset of the strains present in the colony [30,31]. The population is dominated by around twenty species of bacteria. Many of the genera are also found in human gut microbiomes, whilst some are more specialised to insect guts. There is also emerging evidence of their symbiotic role in aiding host metabolism, immunity, and physiology [32].

Here, we test the prediction that higher relatedness should lead to more cooperation using two complementary approaches. First, we follow and extend the approach of Simonet & McNally [19], by testing the prediction that species with higher relatedness have more genes for cooperative traits, or a higher proportion of genes for cooperative traits. We estimate relatedness for each species by comparing genetic similarity within individual bees with genetic similarity across all bees, using the gut metagenomes of 29 worker bees from one colony. We use a new genomic tool SOCfinder [20], that combines information from several methods, to identify genes for cooperative traits. Second, we use molecular population genetics to test the predictions that: (a) genes for cooperative traits favoured by kin selection have increased nucleotide polymorphism and sequence divergence; (b) species with lower relatedness have a stronger signature of kin selection on their genes for cooperative traits. We also use this signature to give a second estimate of relatedness.

## Results

We used previously published data from metagenomic sequencing of 29 *A. mellifera* worker bee guts from a single colony [28]. From these bees, we were able to analyse 19 bacterial ‘sequence discrete populations’ (species) which met minimum requirements for abundance and genomic coverage. For each bacterial species we identified genes for putatively cooperative traits using SOCfinder, used molecular population genetics to assess how selection is acting on those genes, and used metagenomic data to estimate relatedness for each species. We then used phylogenetic linear mixed-effect models, which control for the shared ancestry of species [33], to look for associations between metagenomic relatedness and the number of genes for cooperative traits, and how selection acts on those genes.

### Species vary in how many genes for cooperative traits they carry

We used two established methods to count the number of genes for cooperative traits in each genome. First, we used SOCfinder, which categorises a gene as being for a cooperative trait if it is predicted to code for an extracellular protein, have a cooperative functional annotation, or be part of the biosynthesis of a cooperative secondary metabolite. SOCfinder classed 0.9% to 3.4% of the genes in each genome as being for cooperative traits (Supplement S2). We also used the method of Simonet & McNally, which looks for secreted proteins and genes annotated with cooperative functions, to allow direct comparison between our studies. There is a positive correlation in the number of genes for cooperative traits found by these two methods (Pearson’s r = 0.55, p = 0.01), with SOCfinder finding on average 40 genes for cooperative traits (range: 14–69) compared to 29 by the Simonet method (range: 12–57; Figure S2).

### Species vary in their metagenomic relatedness and relative abundance

We used the metagenomic analysis pipeline tool MIDAS to estimate relative abundance and relatedness for each species. Relatedness is a measure of the genomic similarity of interacting individuals relative to the average genomic similarity in the population. At one extreme, if each bee contains unique variation, then individuals are interacting with social partners more similar to them than the population average, and relatedness is high. At the other extreme, if all of the variation in the population is also present within each bee, then relatedness is low (Figure 1). To estimate relatedness for a species, we therefore compared average genomic similarity within each worker bee with average similarity between bees, using the standard equation for relatedness [34]. We call this measure ‘metagenomic relatedness’. There was substantial variation in the average relatedness of species, from Gilli-1 at 0.38, to Commensalibacter at 0.87 (Figure 2). We also found substantial variation in relative abundance (Figure S3).

**Figure 1:**
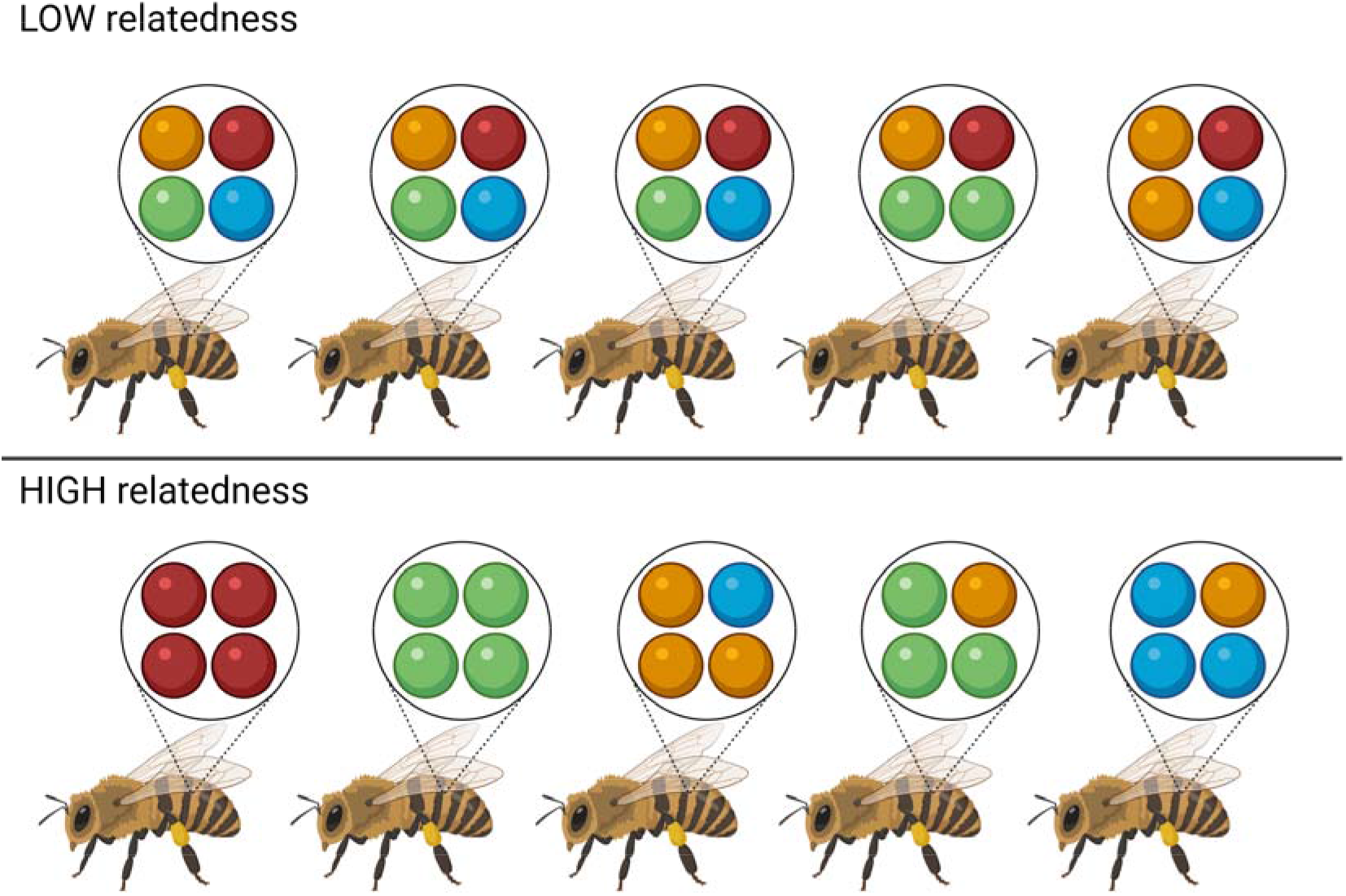
The concept of estimating relatedness from metagenomic data. Within each bee there is genetic variation, represented by the four different colour balls (orange, red, green and blue). We can measure genomic similarity by calculating the average probability of a genomic site being identical in the global population across all bees (*B*), and the average probability of a genomic site being identical within each bee (*W*). Relatedness compares these two measures; *R* = (*W* – *B*) / (1 – *B*). Relatedness is low when the average genomic similarity within each host is close to that expected from the average in the global population (top panel). Relatedness is high when the genomic similarity within a host is higher than expected from the average in the global population (bottom panel). Created in BioRender. Belcher, L. (2027) https://BioRender.com/opalghe

**Figure 2:**
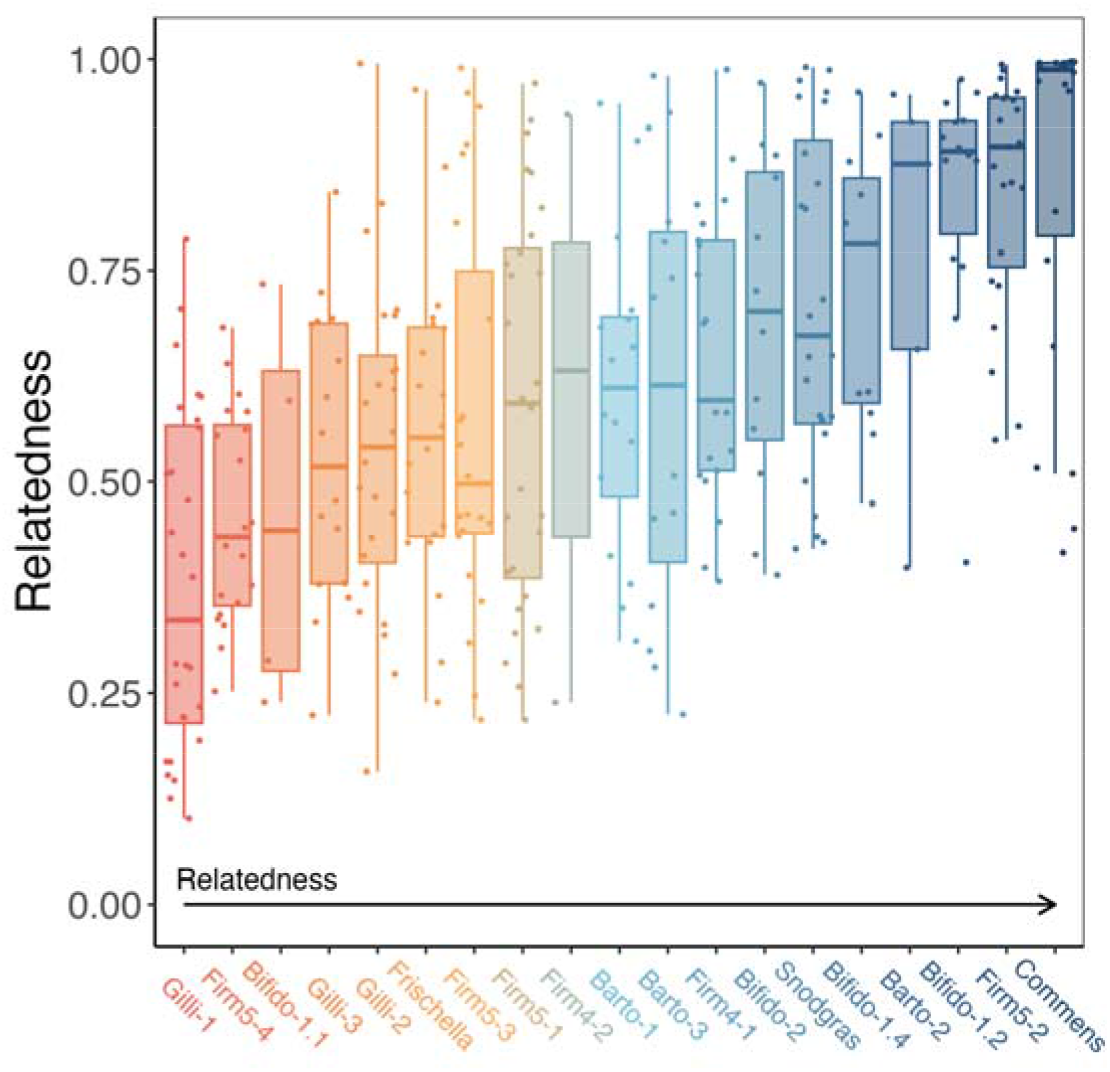
Metagenomic relatedness estimate for 19 species of bacteria in 29 bee metagenomes. Each point represents a point estimate of relatedness in one bee. Species are ordered from lowest to highest relatedness.

### Species show a consistent signature of kin selection

Theory predicts that when relatedness is less than one, selection will be relaxed on genes for cooperative traits relative to those for private traits, because the gene gives a benefit to individuals who do not always carry it. Relaxed selection allows more mutations that change the protein to persist. To test this, we measured nucleotide diversity (polymorphism) in each gene, separately at non-synonymous sites, where a mutation changes the protein, and at synonymous sites, where it does not. We compared genes for cooperative traits with genes for private traits, which benefit only the cell that expresses them. Across the 19 species, genes for cooperative traits had 36% more non-synonymous relative to synonymous polymorphism than genes for private traits (phylogenetic mixed model controlling for gene length; ratio = 1.36, 95% CI = 1.08–1.65, pMCMC = 0.016; higher in 17 of 19 species; Figure 3, Table S10). This suggests that genes for cooperative traits are evolving under kin selection, as they appear to be in other species.

**Figure 3:**
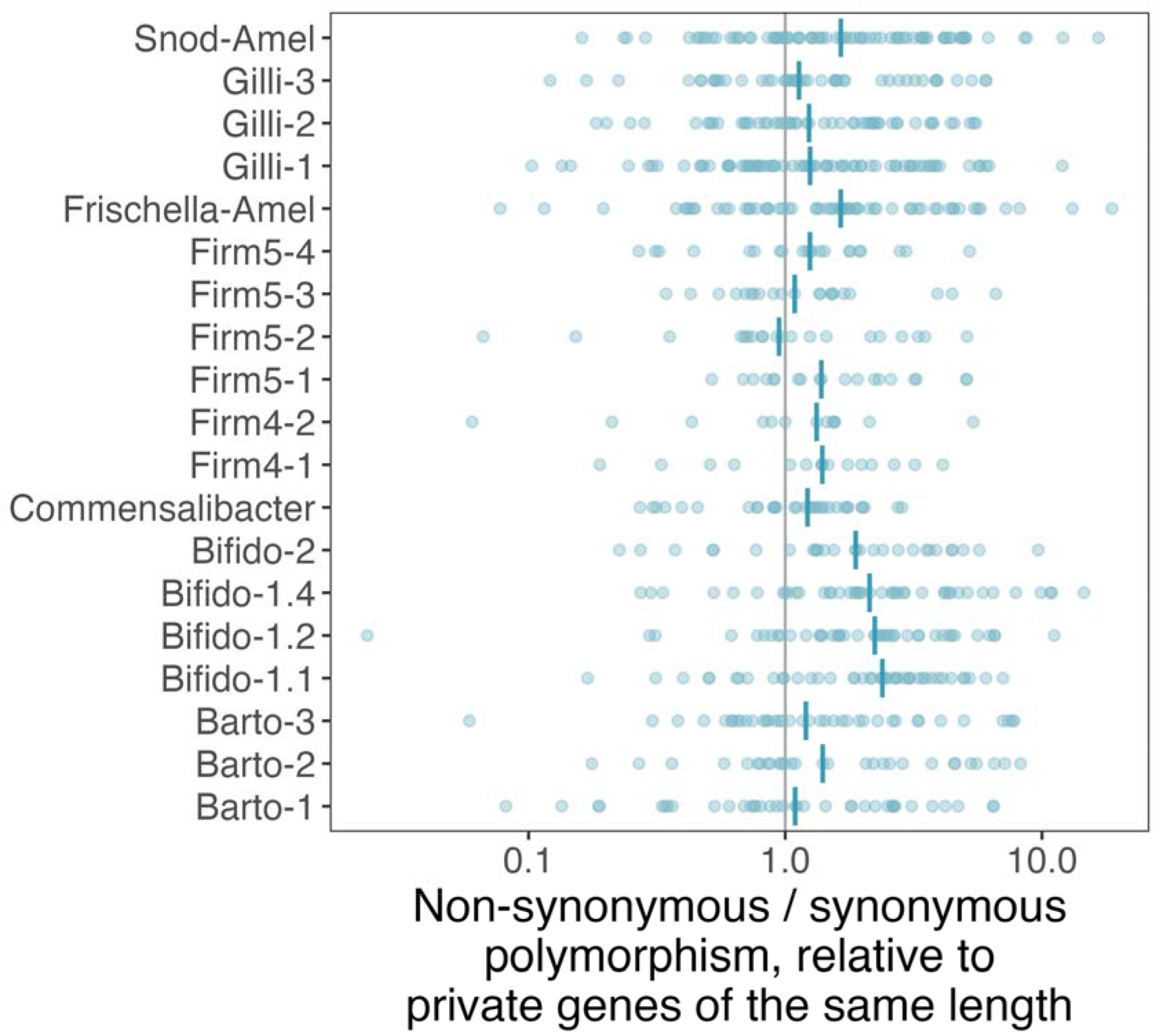
Non-synonymous relative to synonymous polymorphism in genes for cooperative traits, for 19 species of bacteria. Each point is a gene for a cooperative trait. For each gene, we divided its ratio of non-synonymous to synonymous polymorphism by the value expected for a gene for a private trait of the same length in that species. Points to the right of the grey vertical line at 1 have more non-synonymous relative to synonymous polymorphism than genes for private traits. The vertical blue line for each species shows the median. The x-axis is on a log scale.

### Relatedness estimated from polymorphism

The difference between genes for cooperative and private traits also gives a second estimate of relatedness. If selection on genes for cooperative traits is relaxed in proportion to relatedness, their polymorphism is expected to be 1/r times that of genes for private traits (Figure S1C) [27]. The ratio of 1.36 therefore implies an average relatedness of 0.73 between interacting cells (95% CI = 0.61–0.92), similar to the 0.77 estimated in this way for *Bacillus subtilis* [27]. Estimated species by species, this relatedness from polymorphism ranged from 0.47 to 1.09, with a median of 0.77 (Supplement S11). It was higher than the average metagenomic relatedness of 0.63, as expected if bacteria interact in patches within the gut (see Discussion).

### Genes for cooperative traits show the same pattern in three more populations

To test the robustness of our result, we repeated the analysis on metagenomes from three more populations of bees. These came from two colonies in Lausanne, Switzerland, one of which was sampled in two years [35]. To make the test independent of our original methods as well, we used a different pipeline (MIDAS2 with the GTDB database). This pipeline covered 15 of our 19 species, and metagenomic relatedness estimated with it agreed closely with our original estimates (Spearman’s ρ = 0.87, n = 15 species; Supplement S5).

Genes for cooperative traits again had more non-synonymous relative to synonymous polymorphism than genes for private traits. In the three new populations, which are independent of our original data, the ratio was 17% higher in genes for cooperative traits (mixed model controlling for gene length; ratio = 1.17; Figure 4), and it was higher in 34 of 42 cases of a species in a population. The ratio for each species in each population is in Supplement S5 (Figure S5). This ratio implies a relatedness from polymorphism of 0.85 in these populations.

**Figure 4:**
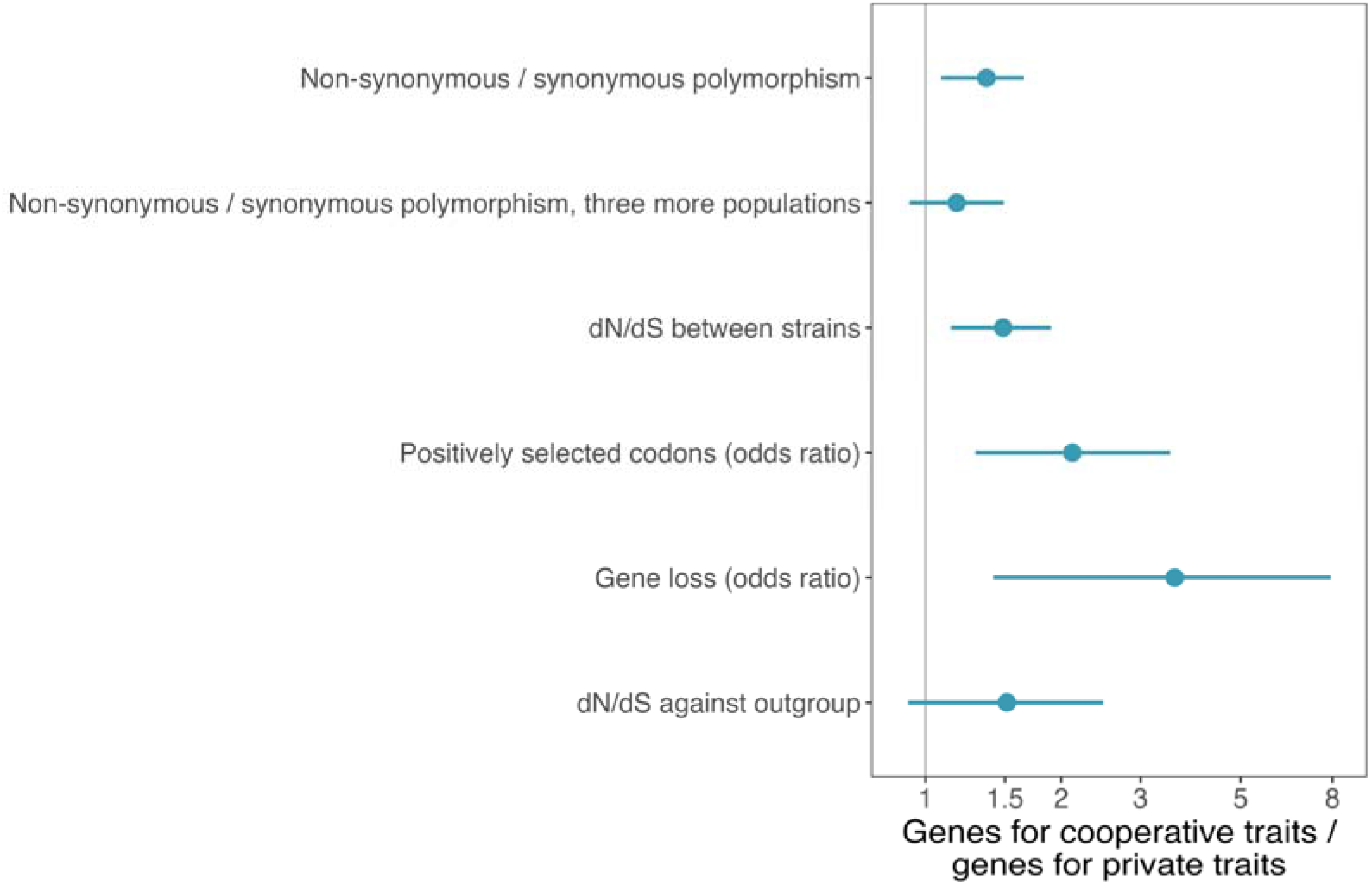
Genes for cooperative traits compared with genes for private traits at each timescale that we measured. Each point is the estimate for genes for cooperative traits relative to genes for private traits, with its 95% credible interval from phylogenetic mixed models. Values to the right of the grey line at 1 mean the measure is higher in genes for cooperative traits. From top to bottom, the rows show non-synonymous relative to synonymous polymorphism within bees in the OH colony (19 species) and in the three new populations (15 species), dN/dS between strains (10 species), the odds that an orthogroup has a positively selected codon (10 species), the odds that an orthogroup has been lost on the strain tree (16 species), and dN/dS between each reference genome and its outgroup (13 species). Non-synonymous relative to synonymous polymorphism and dN/dS are ratios, and positive selection and gene loss are odds ratios, so their sizes cannot be compared directly with the ratios. The x-axis is on a log scale.

### Genes for cooperative traits evolve faster between strains

To look at evolution over a longer timescale than polymorphism within a colony, we compared the genomes of different strains of each species. We grouped each gene with the matching genes in the other strains (an orthogroup), and used orthogroups present in at least eight strain genomes. This gave 353 orthogroups for cooperative traits and 1,023 for private traits from 10 species (Methods).

Genes for cooperative traits built up more protein-changing mutations between strains than genes for private traits, as expected if selection on them is relaxed. The ratio of non-synonymous to synonymous changes (dN/dS) was 49% higher in genes for cooperative traits than in genes for private traits (median 0.075 vs 0.049; mixed model controlling for gene length; ratio = 1.49, 95% CI = 1.14–1.90, pMCMC = 0.008; Figure 4, Table S4).

Genes for cooperative traits were also more likely than genes for private traits to contain codons where selection has favoured changes to the protein (positive selection). Using a codon-level test (FEL), we found at least one such codon in 30% of orthogroups for cooperative traits and 13% of those for private traits (mixed model controlling for gene length, synonymous change and recombination; odds ratio = 2.12, 95% CI = 1.29–3.49, pMCMC = 0.011; Figure 4). We controlled for recombination (exchange of DNA between strains) because it can produce false signals of positive selection. Further checks of these results are in Supplement S6.

### Genes for cooperative traits are lost more often

Genes for cooperative traits were also lost more often than genes for private traits, as expected if selection on them is relaxed. We mapped losses of orthogroups onto the family tree of strains (strain tree) of each of the 16 species with at least five strain genomes. We used GLADE [36], which reconstructs gene losses from gene trees (Methods). Orthogroups for cooperative traits were more likely than those for private traits to have been lost on at least one branch (15% vs 7%; mixed model; odds ratio = 3.57, 95% CI = 1.41–7.93, pMCMC = 0.015; Figure 4). The difference was largest in Bifido-1.1, Firm5-4, Bifido-1.2 and Bifido-1.4 (Table S3). Across all genes, lost orthogroups were enriched for carbohydrate metabolism and transport (Supplement S7).

### Genes for cooperative traits show a trend toward faster divergence from related species

Genes for cooperative traits also showed a trend toward faster divergence from related species than genes for private traits, as expected if selection on them has been relaxed over longer periods. We compared each reference genome with a closely related species (the outgroup; Methods). In 11 of the 19 species the outgroup was so distant that silent changes could not be counted reliably in most genes. We removed genes in which silent changes could not be counted reliably (Methods). This left 237 genes for cooperative traits and 772 for private traits from 18 species, 14 of which had genes for cooperative traits (Table S5). Genes for cooperative traits showed a trend toward higher dN/dS than genes for private traits, but this trend was not significant (median 0.069 vs 0.050; mixed model of the 13 species with at least five genes for cooperative traits, controlling for gene length; ratio = 1.51, 95% CI = 0.92–2.48, pMCMC = 0.085; Figure 4).

### Metagenomic relatedness does not predict the signature of kin selection

Theory predicts that species with lower relatedness should show a larger difference between genes for cooperative and private traits, due to more relaxed selection on genes for cooperative traits. We found no significant correlation between metagenomic relatedness (Figure 2) and the difference in non-synonymous relative to synonymous polymorphism (MCMCglmm; posterior mean = 0.08, 95% CI = −0.63 to 0.87, pMCMC = 0.84; Figure 5).

**Figure 5:**
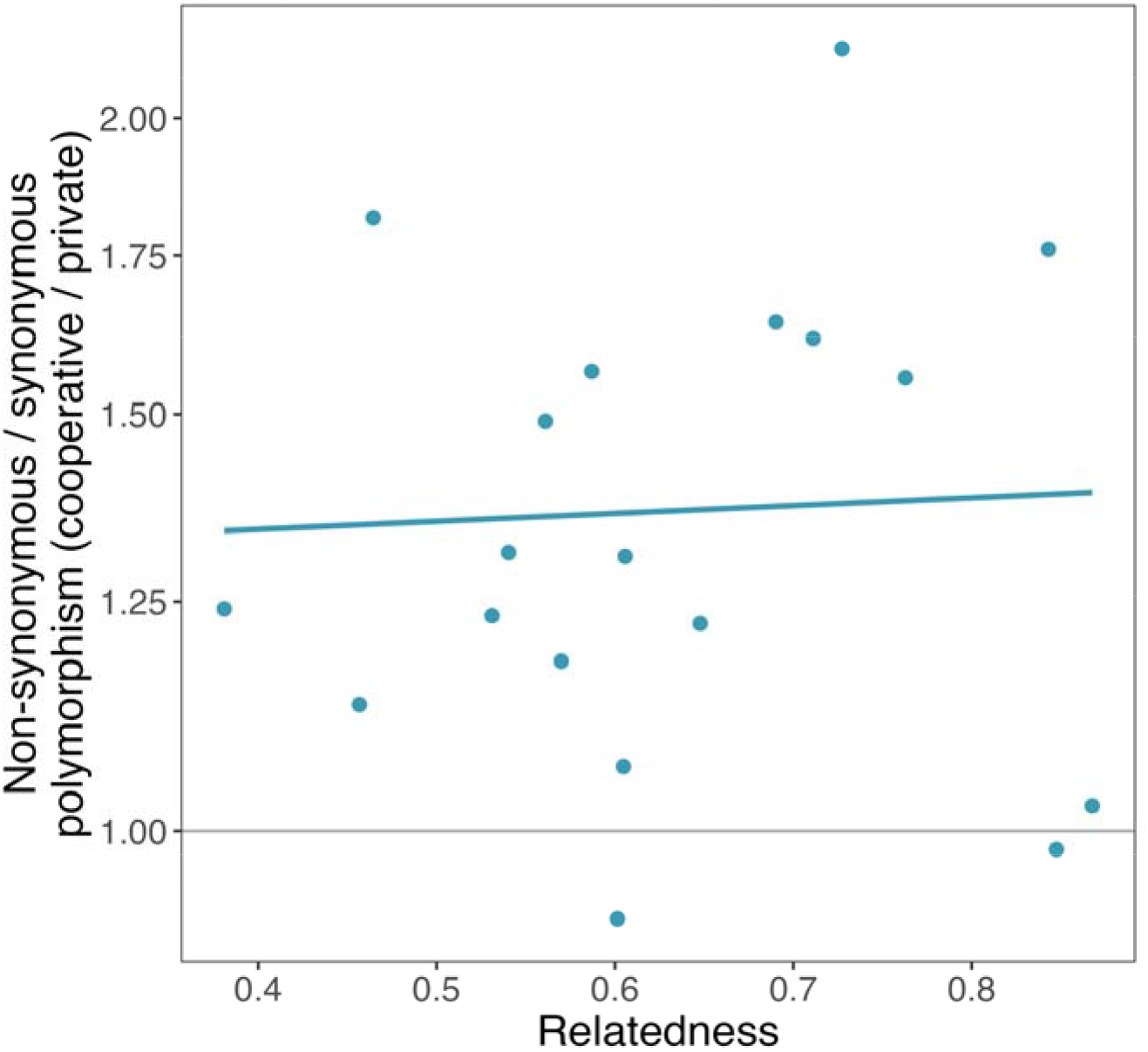
Metagenomic relatedness and the difference in polymorphism between genes for cooperative and private traits. For each species, the y-axis shows the ratio of non-synonymous to synonymous polymorphism in genes for cooperative traits relative to genes for private traits of the same length (log scale). Theory predicts that species with lower relatedness should have a larger ratio, due to more relaxed selection on genes for cooperative traits. The line shows the relationship estimated by the phylogenetic mixed model (posterior mean).

Relatedness as estimated by metagenomics also did not influence how much genes for cooperative traits differed from genes for private traits in non-synonymous relative to synonymous polymorphism across the four populations, dN/dS between strains, positive selection, gene loss or dN/dS against the outgroup (MCMCglmm; pMCMC = 0.52 to 0.98; Supplement S8).

### Metagenomic relatedness does not predict the number of genes for cooperative traits

Species with higher metagenomic relatedness did not have more genes for cooperative traits. Metagenomic relatedness did not predict the number of genes for cooperative traits found by SOCfinder (MCMCglmm; posterior mean = −0.11, 95% CI = −0.99 to 0.73, pMCMC = 0.80), the proportion of genes that were for cooperative traits (posterior mean = −0.11, 95% CI = −0.95 to 0.71, pMCMC = 0.79), or the number found with the method of Simonet & McNally (posterior mean = 0.55, 95% CI = −0.54 to 1.58, pMCMC = 0.30). Relatedness from polymorphism did not significantly predict them either (pMCMC = 0.06 to 0.37; Supplement S11).

### We have reasonable statistical power to detect an effect

We also conducted a phylogenetic power analysis. We simulated data for our species using the effect size reported in Simonet & McNally as the basis for simulating the number of genes for cooperative traits for each value of metagenomic relatedness. We then ran MCMCglmms on this simulated data and found a significant effect of relatedness on the number of genes for cooperative traits in 57% of the simulated runs, implying that we have reasonable statistical power to detect an effect (Supplement S10).

## Discussion

We found consistent evidence for a signature of kin selection on genes for cooperative traits across 19 species of bacteria (Figures 3 and 4). Our measures of kin selection varied across species, both in terms of the number of genes for cooperative traits, and the signatures of selection acting on those genes (Figures 2 to 4). However, in contrast with theoretical predictions, we found no strong evidence that the signature of kin selection or proportion of genes for cooperative traits varies with metagenomic relatedness (Figure 5).

We found that genes for cooperative traits had more non-synonymous relative to synonymous polymorphism than genes for private traits across 19 species of bee gut bacteria, and that this held in three further populations (Figures 3 and 4). Between strains, genes for cooperative traits had higher dN/dS, more positively selected codons and more frequent gene loss, and they showed a trend toward faster divergence from related species (Figure 4). This agreed with our previous analyses of two other species, *Pseudomonas aeruginosa* and *Bacillus subtilis* [14,27]. Taken together, these results provide strong consistent support for the hypothesis that bacteria cooperate, and that this cooperation is favoured by kin selection. Relaxed selection alone would not produce the excess of positively selected codons in genes for cooperative traits. Some of these genes may therefore also be under repeated adaptive change, as might happen if they are involved in conflict between strains [37] or with the host. Genes for cooperative traits are also lost more often than genes for private traits in other bacteria, and in other studies of bee gut bacteria. Genes for cooperative traits are more often present in only some strains of a species (accessory genes), and are lost more often within species than other genes [38]. In bee gut bacteria, secretion systems are lost repeatedly within lineages [39].

In contrast with Simonet & McNally’s study on the human microbiome, we did not find that bee microbiome species with higher metagenomic relatedness had either more or a higher proportion of genes for cooperative traits. This result held when we used their method for counting genes for cooperative traits. There are a number of issues which could have limited the statistical power of our analysis. In particular: (a) we analysed far fewer species - 19 compared to 101 – which our power analysis suggested would have led to reasonable but not high statistical power; (b) many of our species are not well studied and only recently described, possibly limiting our ability to identify genes for cooperative traits; (c) there are potential limitations with our metagenomic measures of relatedness because they assume that bacteria interact across the whole bee gut (see below). However, similar concerns could apply to Simonet & McNally’s study; we did find considerable variation to explain (metagenomic relatedness ranged from 0.38 to 0.87, a spread similar to that in the 101 species of Simonet & McNally); and SOCfinder did not find more genes for cooperative traits in the species that are better studied (Spearman’s ρ = 0.18, p = 0.46, using the proportion of proteins with a known function as a measure of how well studied a genome is; Supplement S2).

We also found no support for our prediction that the difference between genes for cooperative and private traits would be larger in species where metagenomic relatedness is lower (Figure 5). There are at least five possible reasons for this. First, as discussed above, there are potential problems with how we have estimated metagenomic relatedness, because we have had to assume that all the bacteria within each host are equally able to interact with each other. When this is not the case, and bacterial interactions are more local within the host gut, then we would have underestimated relatedness. Consequently, our metagenomic estimate of relatedness is better seen as a lower bound, such that relatedness is between the estimated value and 1. For example, strains of *Snodgrassella* can coexist in one bee by occupying separate patches of the ileum in the hindgut [40]. The strains that colonise a bee first tend to dominate it [41], and so local patches of related cells may be common, leading to relatedness being higher than our metagenomic estimate. Consistent with this, relatedness estimated from polymorphism (0.73) was higher than the average metagenomic relatedness (0.63). Second, our ability to identify genes for cooperative traits may not have been accurate enough, in these relatively little studied species [20]. There is also the complication that some behaviours may involve cooperation with the host rather than their own species [42]. Third, the predicted influence of relatedness assumes that ‘all else is equal’ [21,22,26]. If there are other factors that vary across species, such as the relative costs and benefits of different behaviours, which we cannot control for, then this would add noise that would prevent the prediction from being observed. Fourth, we might not have been able to measure the signature of selection accurately enough from metagenome data, particularly on species which are at relatively low abundance within each bee gut. Species at low abundance contribute fewer reads in each bee, so fewer sites can be measured and estimates of polymorphism are noisier. Fifth, some bacterial behaviours may involve cross-feeding - interactions that have not been explicitly modelled with theory and can alter how selection acts on the genes controlling those traits. If species can cross feed on each other’s metabolites [43,44], or utilise acids provided by the bee [45], then selection on genes involved in those traits will change.

There are at least two possible limitations and complicating factors with our population genetic analyses. First, we used an automated method (SOCfinder) to identify genes for cooperative traits. Whilst this method facilitates broad comparative studies based on many genes, it is likely to be less accurate than the artisanal annotation of genes based on data from laboratory experiments [14,20,27]. Our previous tests of SOCfinder have shown that it detects signatures of kin selection in the same direction as those based on artisanal annotation, but it is possible that extra noise is added, making it harder to examine variation in the level of signature [20]. Second, there are several other factors that can lead to the extent of relaxed selection varying across species [11,23,26,46,47]. The costs and benefits of cooperation may differ between species, for example with diet, cell density or the other species present. Another possibility is conditional expression. If a gene only has effects in certain environments, or in certain generations, then this leads to a similar signature of selection as kin selection.

To conclude, both phenotypic and population genetic analyses have found consistent evidence for cooperation favoured by kin selection in natural populations of bacteria [14,17,19,27,42]. A major challenge for the future is to determine whether variation in the level or form of cooperation across species can be explained by kin selection theory, analogous to the patterns that have been observed in animals [18]. While relatedness is predicted to play a central role, there are many factors that can influence the cost and benefit of cooperation, and hence how selection acts on the genes controlling cooperative traits.

## Methods

### Metagenomes

We used 29 metagenomes from individual honeybees in a colony of the OH subspecies (*Apis mellifera mellifera*) sequenced and published by Wu *et al*. [28].

### Metagenomic analysis

We followed the method outlined in Simonet & McNally [19] of using the established metagenomic pipeline MIDAS [48] to estimate relative abundance and relatedness for each species. We briefly explain the relevant methods in each section below.

### Building a custom MIDAS database

The MIDAS pipeline involves mapping metagenomic sequences to a database of genomes, in order to profile which species are present. Because the default database misses many bee microbiome species, we built a custom MIDAS database of sequenced genomes of bee gut bacteria using the build_midas_db.py script provided by MIDAS. We used the list of 405 sequenced genomes in the supplement of Wu *et al.* [28] as a starting point, and filtered out those which did not exist in the PATRIC database, or did not match a known bee microbiome species. This led to n=334 genomes from n=35 ‘sequence discrete populations’ (SDPs). A custom MIDAS database requires a representative (reference) genome for each SDP, which we selected manually by focusing on maximising completeness of genome and minimising number of contigs. We then ran MIDAS for our 29 metagenomes using this custom database. We used default minimum coverage requirements.

We used the script for calculating relatedness from a MIDAS output provided by Simonet & McNally [19]. Briefly, we can measure genomic similarity by calculating the average probability of a genomic site being identical in the global population across all bees (*B*), and the average probability of a genomic site being identical within each bee (*W*). Relatedness compares these two measures [19,34]; *R* = (*W* – *B*) / (1 – *B*). We need at least two hosts to measure relatedness, so we filtered out those SDPs only present in one host.

### Relative abundance

Relative abundance was estimated from the number of sequence reads that map to the reference genome for each species, and is extracted directly from the MIDAS output.

### Bacterial species

We analysed 19 bacterial SDPs which met minimum requirements for abundance, genomic depth, coverage, and presence in multiple hosts. SDPs are a method of splitting genomes into evolutionary units based on gaps in the distribution of pairwise average nucleotide identity [49,50]. These boundaries exist across prokaryotes, including in honey bee microbiome bacteria. In the bee gut, strains of the same *Lactobacillus* species compete more strongly than strains of different species, which supports treating these species as ecological units [51]. We used this categorisation of genomes into SDPs from Ellegaard & Engel [35]. For simplicity, we refer to each SDP as a species. The 19 species we analysed capture all of the known dominant representatives of the gut, and are listed in Table 1.

**Table 1:** List of species used in the analysis.

| **SDP name** | **Genus** | **Reference Accession** |
|---|---|---|
| **Barto-1** | *Bartonella* | GCA_016100345.1 |
| **Barto-2** | *Bartonella* | GCA_002007565.1 |
| **Barto-3** | *Bartonella* | GCA_002007505.1 |
| **Bifido-1.1** | *Bifidobacterium* | GCA_016101605.1 |
| **Bifido-1.2** | *Bifidobacterium* | GCA_016100725.1 |
| **Bifido-1.4** | *Bifidobacterium* | GCA_007559155.1 |
| **Bifido-2** | *Bifidobacterium* | GCA_000737865.1 |
| **Commensalibacter** | *Commensalibacter* | GCA_016101865.1 |
| **Firm4-1** | *Bombilactobacillus* | GCA_000967245.1 |
| **Firm4-2** | *Bombilactobacillus* | GCA_016102125.1 |
| **Firm5-1** | *Lactobacillus* | GCA_003150935.1 |
| **Firm5-2** | *Lactobacillus* | GCA_000761135.1 |
| **Firm5-3** | *Lactobacillus* | GCA_003202825.1 |
| **Firm5-4** | *Lactobacillus* | GCA_003151025.1 |
| **Frischella-Amel** | *Frischella* | GCA_000807275.1 |
| **Gilli-1** | *Gilliamella* | GCF_001693435.1 |
| **Gilli-2** | *Gilliamella* | GCA_016101145.1 |
| **Gilli-3** | *Gilliamella* | GCA_007559165.1 |
| **Snod-Amel** | *Snodgrassella* | GCA_002777735.1 |

### Counting genes for cooperative traits

Cooperative traits provide a benefit to many cells within a group, not just those who produce the trait. We used several established methods to count the number of genes for cooperative traits in each genome. First, we used the method outlined in Simonet & McNally [19] which looks for secreted proteins and genes annotated with cooperative functions. This allows us to make a direct comparison between our study and theirs. We also used SOCfinder [20], which is a newer method that builds on the Simonet & McNally [19] method, and categorises a gene as being for a cooperative trait if it is predicted to code for an extracellular protein, have a cooperative functional annotation, or be part of the biosynthesis of a cooperative secondary metabolite. SOCfinder is more effective at finding known genes for cooperative traits, so we use this method for our main analysis. To count the genes for cooperative traits in each species, we applied both methods to two to five genomes of the species and took the median. We compared genes for cooperative traits with genes for private traits whose proteins are predicted by PSORTb [v3.0, 52] to stay in the cytoplasm, and so are the least likely to have a cooperative function, as in our previous work [14,20,27].

Methods for counting weapons [53] and carbohydrate-active enzymes are in Supplement S9.

### SNP-calling

We used an established SNP-calling pipeline to find the SNPs where the sequence in each individual differs from the reference genome for that species. The full details can be found in Belcher *et al.* [14,27], but we explain briefly here. We used Trimmomatic [54] to remove low quality reads. Next, we mapped reads to the relevant reference genome using BWA [55]. We converted the resulting SAM files to sorted BAM files using SAMtools [56]. Then, we called variants using BCFtools [57], and converted the output to a VCF file for easy analysis. We used the SAMtools vcfutils.pl script to filter for mapping quality and read depth. We also used the coverageBed tool from BEDtools [58] to measure what proportion of each gene’s length had been mapped, so that we could later adjust population genetic measures to the mapped length of the gene rather than the length of the gene in the reference genome.

### Molecular population genetics

To test the prediction that species with lower relatedness have a stronger signature of kin selection on their genes for cooperative traits, we measured molecular population genetic parameters such as nucleotide polymorphism using an established method. The full details can be found in Belcher *et al.* [14,27], but we explain briefly here. We conducted most of the analysis in the PopGenome package in R [59]. Specifically, we calculated nucleotide diversity [60]. Where necessary to obtain per-site measures we calculated separately for synonymous and non-synonymous sites, and scaled to the relevant mapped length. For each gene we then took the ratio of non-synonymous to synonymous polymorphism, using genes with at least some polymorphism of each kind.

### Strain genomes, orthogroups and gene trees

To compare genes for cooperative and private traits across strains of the same species, we grouped the genes of several genomes per species into orthogroups. For each species we used the reference genome plus up to 14 further genomes of the same species, from the genomes listed by Wu et al. [28] that we had downloaded from PATRIC. Where more than 14 were available (four species), we chose those with the highest completeness and then the fewest contigs. Three species had fewer than five genomes (Barto-2, Firm4-2 and Frischella-Amel), so we left them out of the comparisons between strains.

We also needed an outgroup for each species, to root the strain tree and to measure divergence. We chose as the outgroup the closest species that was still clearly distinct. We required candidates to share reciprocal best hits [DIAMOND, 61] with at least 40% of the reference proteins, at a median identity of at most 98%, and chose the candidate with the highest median identity. Several outgroups are themselves among our 19 species (for example, Gilli-3 is the outgroup for Gilli-2).

We ran OrthoFinder [v3.1.5, default settings, 62] twice on the genomes of each species and its outgroup. From the single-copy orthogroups of the first run, we built a strain tree [concatenated protein alignment of up to 150 orthogroups; MAFFT v7.526, 63; IQ-TREE v3.1.3 with the LG+G4 model, 64]. We rooted this tree on the outgroup. In the second run, we used the rooted tree to infer gene trees from multiple sequence alignments.

We classed each orthogroup as being for a cooperative or a private trait according to the SOCfinder label of its gene in the reference genome. Orthogroups for private traits were those whose genes in the reference genome were all predicted to be cytoplasmic. We excluded orthogroups whose reference genes had both cooperative and private labels, and orthogroups with no gene in the reference genome.

### Gene loss

If selection on genes for cooperative traits is relaxed, these genes should be lost more often between strains [38]. To test this, we reconstructed the gains and losses of each orthogroup on the rooted strain tree of each species with GLADE [v1.0.0, 36], which uses the OrthoFinder gene trees. GLADE infers two kinds of loss. In the first, a lineage lacks an orthogroup that is present in its sister lineage. In the second, a lineage has lost one copy after a gene duplication. We counted both kinds of loss for each orthogroup on the strain tree, and recorded whether the orthogroup had been lost at least once.

### Selection between strains

For each orthogroup we estimated how fast its protein changed between strains, and whether particular codons had been under positive selection. We used orthogroups with one copy per genome that were present in the reference genome. In our main analysis we used orthogroups present in at least eight genomes, counting the reference (10 species). To keep run times manageable, we tested every orthogroup for a cooperative trait in each species but only a random sample of 200 of those for private traits that passed these filters (fixed random seed). Of the sampled orthogroups for private traits, we used those whose genes in the reference genome were all predicted to be cytoplasmic.

For each orthogroup we aligned the proteins with MAFFT [v7.526, 63], converted the alignment to codons with pal2nal [v14.1, 65], and built a tree from the codon alignment with FastTree [v2.2.0, GTR model, 66]. We estimated the ratio of non-synonymous to synonymous change (dN/dS) across the whole gene with codeml [model M0, F3×4 codon frequencies, universal genetic code, branch lengths estimated; PAML v4.10.10, 67]. Where gene trees had nodes with more than two branches, which arise when strains are nearly identical, we resolved these nodes into two-way splits with zero-length branches before running codeml (Supplement S6). We excluded orthogroups with almost no synonymous change (synonymous tree length below 10) or with dN/dS of 10 or more.

We detected codons under positive selection with HyPhy [v2.5.103, 68]. We used FEL [69], which detects selection that persists across the tree. We scored an orthogroup as positively selected if it had at least one codon with p ≤ 0.1, with no correction for the number of codons tested.

Recombination between strains can create false signals of positive selection. We therefore tested each alignment for recombination with GARD [HyPhy v2.5.103; nucleotide model, general discrete rate variation, 70], and called an orthogroup recombinant if a model with at least one breakpoint fitted better by AICc. We included recombination as a covariate in our models of positive selection.

### Divergence from the outgroup

To measure protein evolution over a longer timescale, we compared the reference genome of each species with its outgroup. For each orthogroup with one copy in the reference and the outgroup, we estimated dN, dS and dN/dS between the two genomes with yn00 [PAML v4.10.10, 71]. When genomes have diverged too far, synonymous sites become saturated and dS is underestimated. We therefore used only genes with 0.01 < dS < 1, which excluded most genes in the most distant pairs.

### Replication with a second pipeline and further colonies

To test whether genes for cooperative traits show the same pattern in other colonies and with a different pipeline, we repeated the analysis of relatedness and polymorphism in the OH colony and in three populations from a second study. We used the 29 bees from the OH colony [28] and 54 bees from two neighbouring colonies at the University of Lausanne, Switzerland [35; SRA study SRP150166]. Ellegaard & Engel sampled 18 bees from the colony ‘Les Droites’ in 2015, and 18 bees each from ‘Les Droites’ and ‘Le Grammont’ in 2016. We treated each colony-year as a separate population. Ellegaard & Engel sampled bees of mixed ages, and strain composition differs between nurses and foragers [72]. Mixed ages may add noise to our estimates of relatedness, but should matter less for the comparison of genes for cooperative and private traits, which we made within the same samples.

We processed all 83 metagenomes with MIDAS2 [v1.0.0, 73] and its GTDB database [release 202, 74]. We first profiled species in each bee, and kept species found in at least two bees. We then built read pileups against the representative genome of each species, for bees in which that species had a median marker coverage of at least 2 and at least half of its markers covered. We matched our species to MIDAS2 species, and transferred our labels of genes for cooperative and private traits, as described in Supplement S5.

From the pileups we calculated nucleotide diversity (π) at each site as the probability that two reads differ. We used sites with a depth of at least 5 and at most three times the median, present in at least half of the bees. We calculated diversity within each bee and in all bees of a population pooled. We calculated relatedness in each bee as R = 1 − π(within) / π(pooled), equivalent to R = (W − B) / (1 − B), where W = 1 − π(within) and B = 1 − π(pooled). We averaged R over bees, as in our main analysis.

For each gene, we used the read counts pooled over the bees of a population, at sites covered in at least two bees, and used genes with at least half their length covered. We classed each polymorphic site as non-synonymous or synonymous by placing the most common base that differed from the reference into the reference codon, and took the ratio of the summed π at non-synonymous sites to that at synonymous sites. We kept each species in each population if it had at least five bees and at least five genes for cooperative traits (57 cases).

### Statistical modelling

We used Bayesian mixed-effects models implemented in the R package MCMCglmm [75] to compare genes for cooperative and private traits, and to analyse the relationship between our measures of relatedness and our measures of kin selection. This approach is commonly used to control for the phylogenetic relationship between species, which is necessary as species share evolutionary history and are not independent data points [33]. For all models, we included phylogeny as a random effect and report the pMCMC value, which can be interpreted in the same way that we interpret a traditional p-value.

### Statistical models for comparisons between genes

We compared genes for cooperative and private traits by first estimating the difference between them within each species. For polymorphism, dN/dS between strains and dN/dS against the outgroup, we fitted a linear model of the log of each gene’s value, with gene type (cooperative or private) and log gene length as predictors. For positive selection, we fitted a logistic model of whether an orthogroup had a positively selected codon, with gene type, log gene length, the log of synonymous tree length and recombination as predictors. Longer genes give more power to detect selected codons and recombination, and more synonymous change gives more power to detect selected codons. We used a Jeffreys-prior penalty, so that species with few positively selected orthogroups still gave a finite estimate [76]. For gene loss, we took the log odds ratio of loss for orthogroups for cooperative traits relative to those for private traits, adding 0.5 to each cell of the table because some species had no lost orthogroups for cooperative traits. For divergence from the outgroup, we used the 13 species with at least five genes for cooperative traits. We then analysed these differences, one per species, in a phylogenetic mixed model with phylogeny as a random effect and the sampling variance of each difference as known measurement error. The intercept of this model is the average difference between genes for cooperative and private traits, controlling for phylogeny. For dN/dS between strains and positive selection, we also fitted models with every orthogroup as a data point, and phylogeny and species as random effects (Supplement S6).

To ask whether differences between genes for cooperative and private traits varied with relatedness across species, we added metagenomic relatedness as a predictor to these phylogenetic mixed models [75,77] of one value per species. We modelled the number of genes for cooperative traits with a Poisson model, with the log number of other genes in the genome as a covariate, and their proportion with a binomial model, each with metagenomic relatedness as a predictor. For the main models, we used inverse-Wishart priors (V = 1, ν = 0.002) and ran four chains of 1.1 × 10 iterations, with a burn-in of 10 and thinning of 1,000. We checked that the chains had converged (Gelman–Rubin statistic below 1.01, effective sample size at least 3,000 and autocorrelation between successive samples below 0.1). We repeated the main models with a second phylogeny (Supplement S6). The results of all models are in Table S7. We estimated relatedness from polymorphism as 1/ratio, using the ratio and its 95% CI from the main model, and for each species from its own ratio. We then repeated the models of the number and proportion of genes for cooperative traits with relatedness from polymorphism as the predictor (Supplement S11).

In the replication, each species appeared in up to four populations. We therefore estimated the difference between genes for cooperative and private traits for each species in each population, as above. To test whether genes for cooperative traits showed the same pattern in the three new populations, we modelled these differences with phylogeny, species and population as random effects. To test relatedness, we modelled these differences in all four populations in the same way, with relatedness in that population (from MIDAS2) as a predictor.

### Phylogeny

We built a phylogeny of our 19 species from their reference genomes (Figure S4). We ran OrthoFinder [v3.2, 62] on the reference proteomes, and inferred a maximum-likelihood tree from its concatenated alignment of single-copy orthogroups with IQ-TREE [v3.1.4, 64]. We chose the substitution model with ModelFinder among LG and JTT models by BIC [LG+F+R5, 78], and assessed support with 1,000 ultrafast bootstraps. We rooted the tree where OrthoFinder roots its species tree, between the Proteobacteria and the other species, and made it ultrametric by penalised likelihood [λ = 1 in chronos, 79]. All nodes had 100% bootstrap support except one (85%), and the tree had the same topology as the OrthoFinder species tree. As a second phylogeny, we pruned the GTDB bacterial tree to our species [release 202, 74], which also had the same topology, and repeated our main models with it (Supplement S6).

## Data accessibility

All sequence data we analysed are public. The authors confirm that all supporting data, code and protocols have been provided within the article, through supplementary data files, or in the Figshare archive below.

1. The metagenomes of 29 individual bees from the OH colony [28] are in NCBI BioProject PRJNA645015.
2. The metagenomes of 54 individual bees from the colonies ‘Les Droites’ and ‘Le Grammont’ [35] are in NCBI SRA study SRP150166.
3. The accessions of the reference, strain and outgroup genomes for each species are in Table S1.
4. The code and processed data are archived on Figshare (doi:10.6084/m9.figshare.34955158). The archive holds the scripts for annotation, orthology, gene loss, selection, divergence, the replication and the statistical models, and the result tables for each gene.
5. Supplementary Tables S1 to S10 are in one Excel file, and the supplementary text, figures and Tables S11 to S13 are in one further file.

## Acknowledgements

We thank the UNIQ+ research internship programme at the University of Oxford, which supported the internships of Sarah Flint and Harvey Jeffrey. We thank Waldan Kwong for helpful discussions.

## Author contributions

Conceptualization: L.J.B., A.E.D., Z.K., S.A.W. Methodology: L.J.B., S.A.W. Software: L.J.B. Formal analysis: L.J.B., R.R., S.G.F., H.J.J. Investigation: L.J.B., R.R., S.G.F., H.J.J. Data curation: L.J.B. Visualization: L.J.B. Writing, original draft: L.J.B. Writing, review and editing: all authors. Supervision: L.J.B., S.A.W. Funding acquisition: S.A.W.

