## Supplementary Info for "Kin selection for cooperation in the honey bee microbiome"

Tables S1 to S10 are in the accompanying Excel workbook (bee_microbiome_Supplementary_Tables.xlsx), and Tables S11 to S13 are below. Code and data are on Figshare (doi:10.6084/m9.figshare.34955158).

### Supplement S1: Theoretical predictions

Kin selection theory predicts that genes for cooperative traits are under weaker selection than genes for private traits when relatedness is below one, because the benefits of cooperation sometimes go to individuals who do not carry the gene. Figure S1 summarises the predictions that we test.

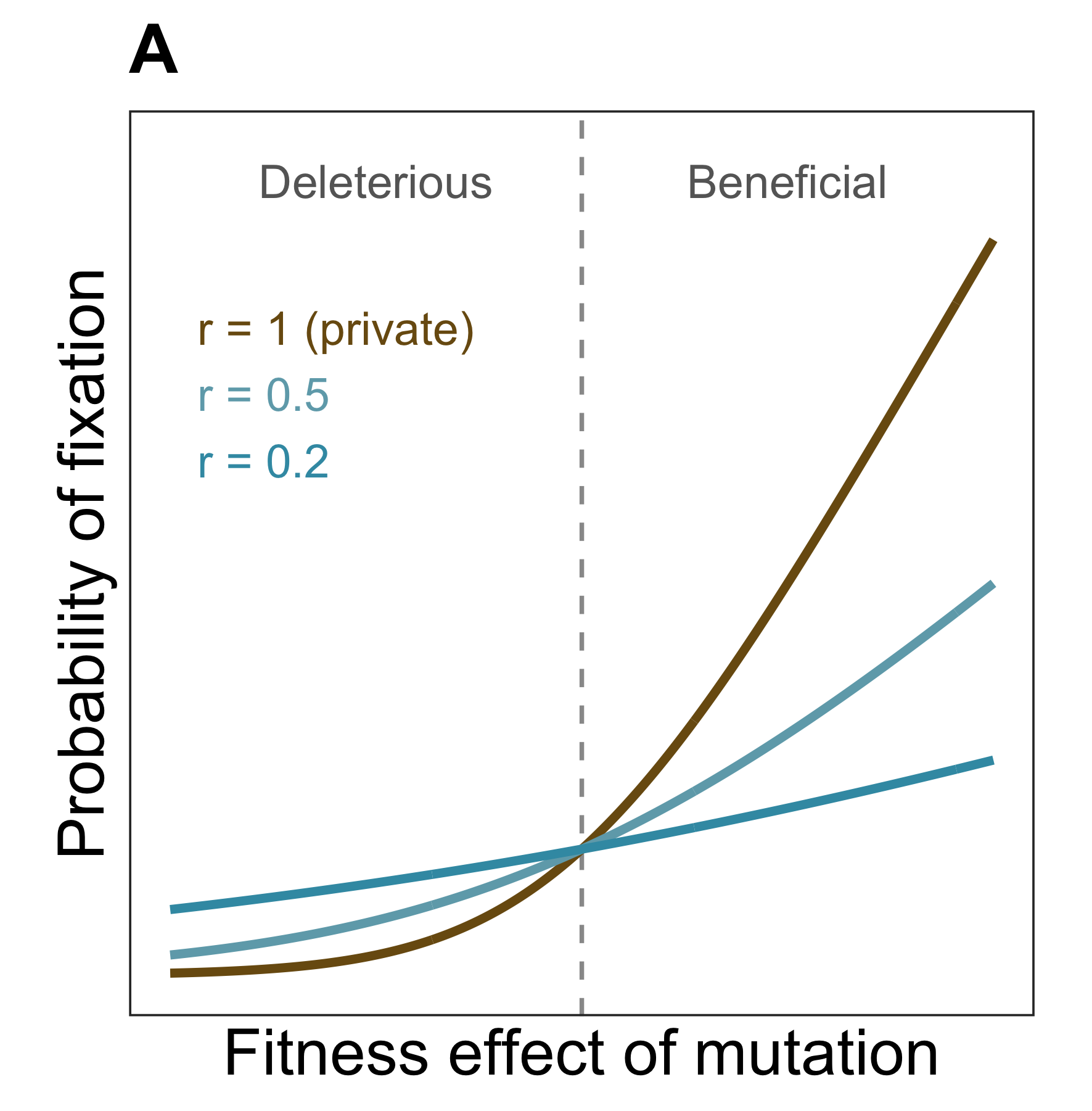

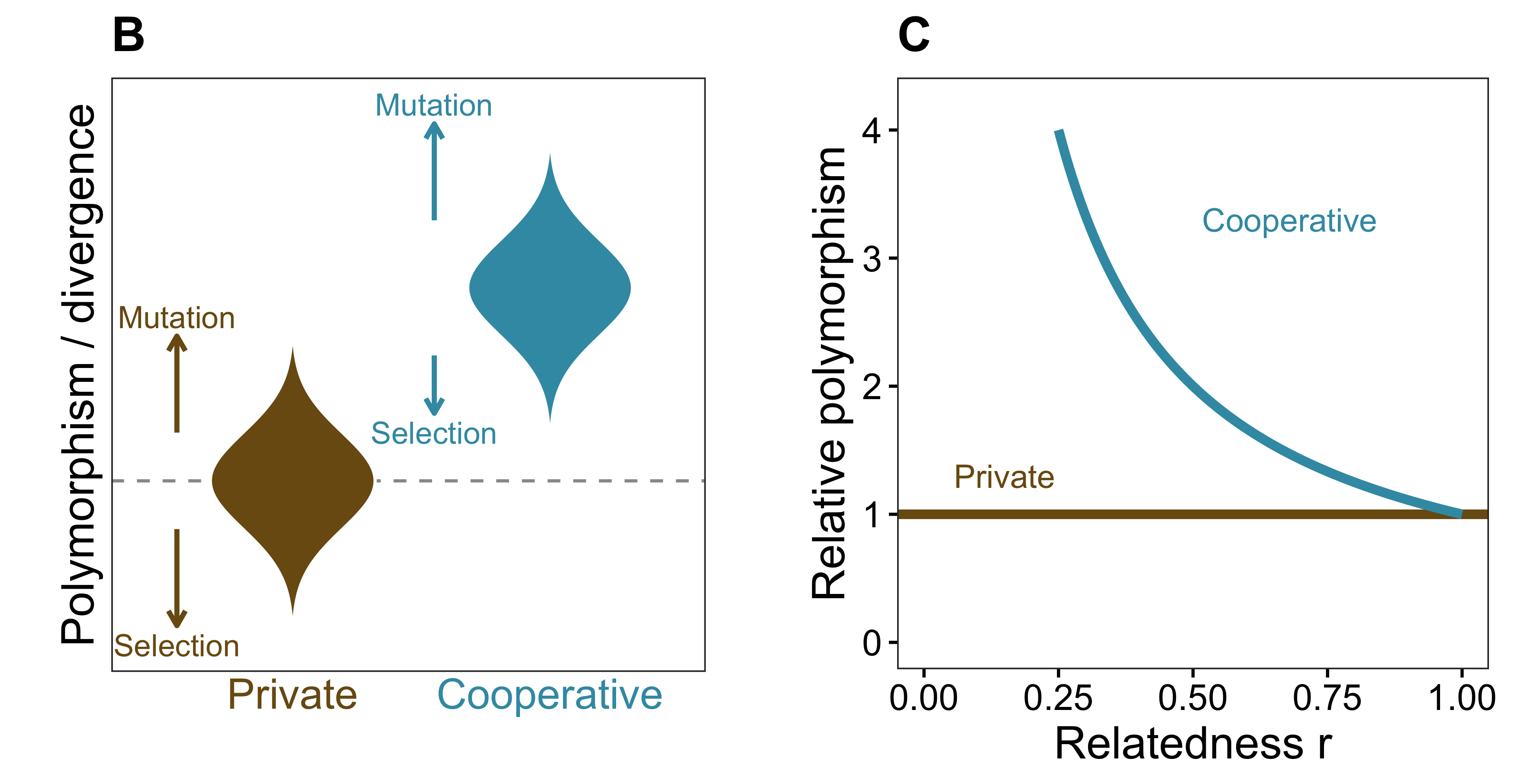

Figure S1: How kin selection shapes the evolution of genes for cooperative traits. (A) The probability that a new mutation becomes fixed, against its effect on fitness, for different values of relatedness. When relatedness is lower, selection is weaker, so harmful mutations are removed less efficiently and beneficial ones are fixed less often. (B) Genes for cooperative traits are therefore expected to have higher polymorphism and divergence than genes for private traits. (C) The expected polymorphism of genes for cooperative traits relative to genes for private traits increases as relatedness falls.

### Supplement S2: Counting genes for cooperative traits

SOCfinder identified 14 to 69 genes for cooperative traits per genome (median over two to five genomes of each species; mean 40), which is 0.9% to 3.4% of all genes. The method of Simonet & McNally (2021) identified 12 to 57 (mean 29). The two counts were positively correlated across species (Pearson’s r = 0.55, p = 0.01; Figure S2).

Species whose genomes are better studied did not have more genes for cooperative traits. We used the proportion of proteins in each reference genome with a known function (those that eggNOG-mapper assigned to a COG category other than ‘function unknown’; 56% to 75% across species) as a measure of how well studied a genome is. This proportion did not predict the proportion of genes that SOCfinder classed as being for cooperative traits (Spearman’s ρ = 0.18, p = 0.46) or their number (ρ = −0.06, p = 0.82).

Metagenomic relatedness varied across our species about as much as across the human gut species of Simonet & McNally (2021). Relatedness ranged from 0.38 to 0.87 in our 19 species (standard deviation 0.14), and from 0.12 to 0.93 in their 101 species (standard deviation 0.15). The main difference in statistical power between the two studies is therefore the number of species.

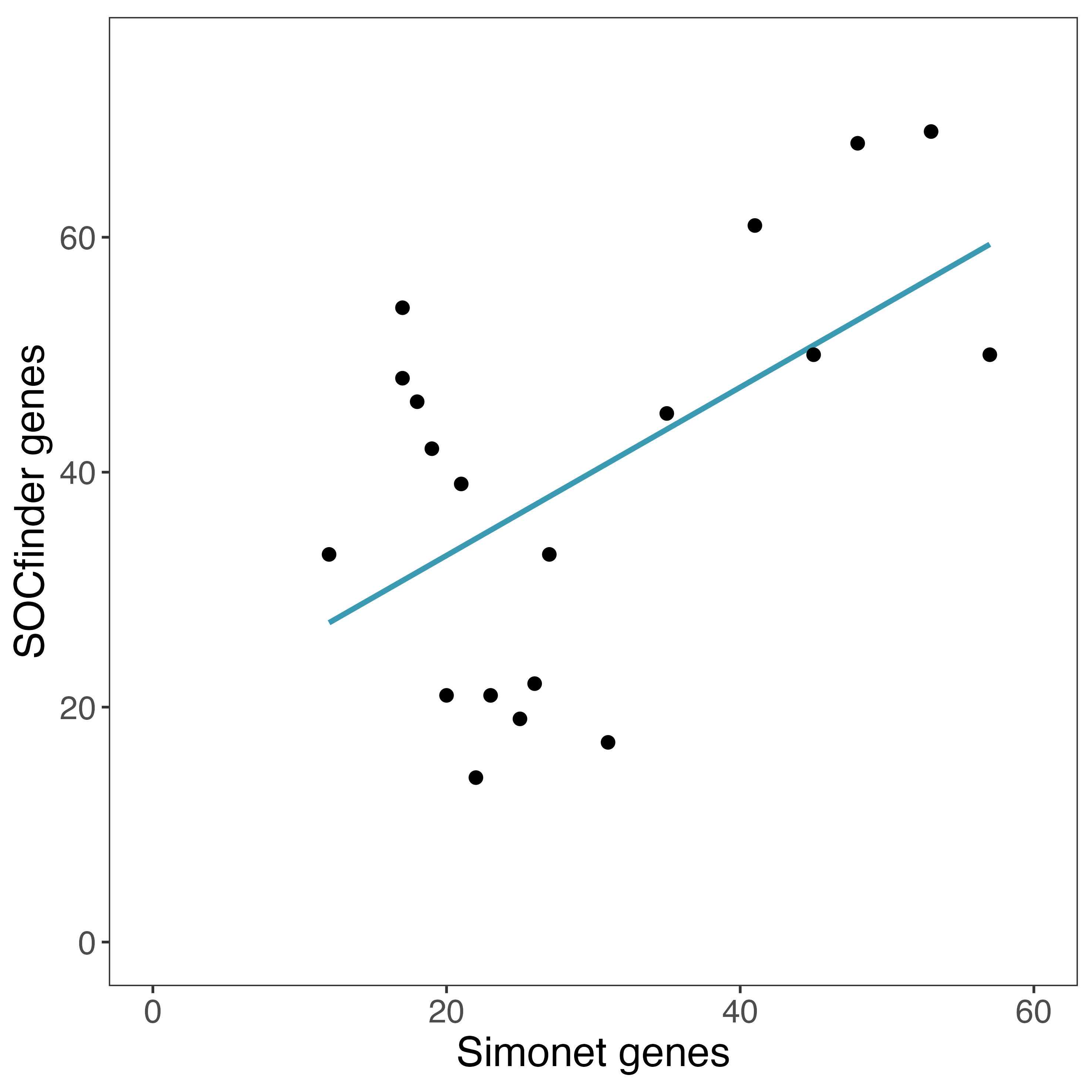

Figure S2: Correlation between the number of genes for cooperative traits found by SOCfinder and by the method of Simonet & McNally (2021).

### Supplement S3: Relative abundance

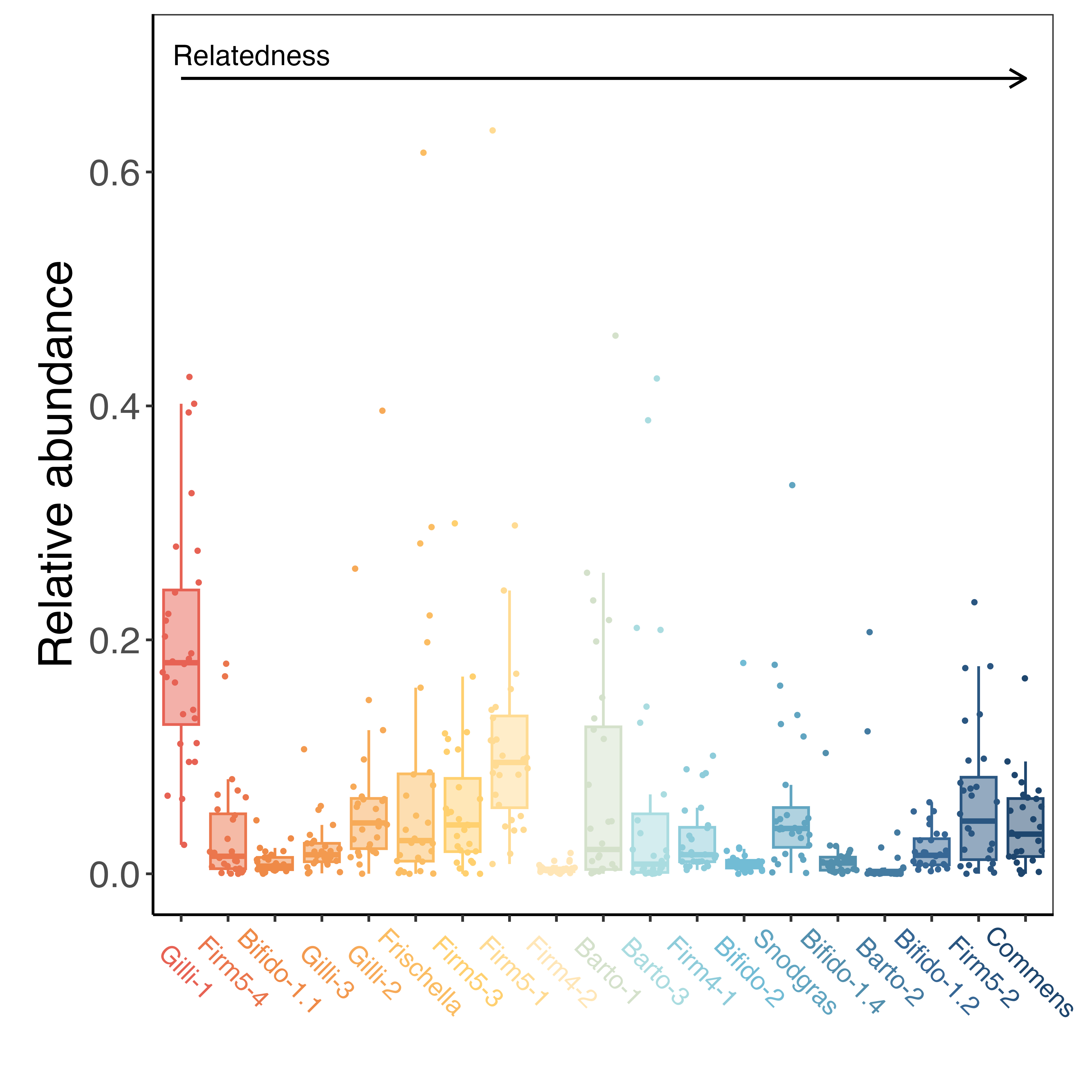

Figure S3: Relative abundance for 19 species of bacteria in 29 bee metagenomes. Each point is the relative abundance in one bee. Species are ordered from lowest to highest metagenomic relatedness.

### Supplement S4: Phylogeny

We built the phylogeny of our 19 species from their reference genomes with OrthoFinder and IQ-TREE, as described in the main Methods. All nodes had 100% ultrafast bootstrap support except the node joining Bifido-1.1 and Bifido-1.2 (85%). The tree had the same topology as the species tree from OrthoFinder.

As a second phylogeny, we pruned the GTDB bacterial tree (release 202; Parks et al., 2022) to our species. For 13 species we used the species’ own genome in GTDB, and for the other six, whose genomes were added after release 202, we used the nearest GTDB representative genome. We rooted this tree between the Proteobacteria and the other species and made it ultrametric by penalised likelihood, as for the main tree. It had the same topology as the main tree. All results with this tree are in Supplement S6 and S8.

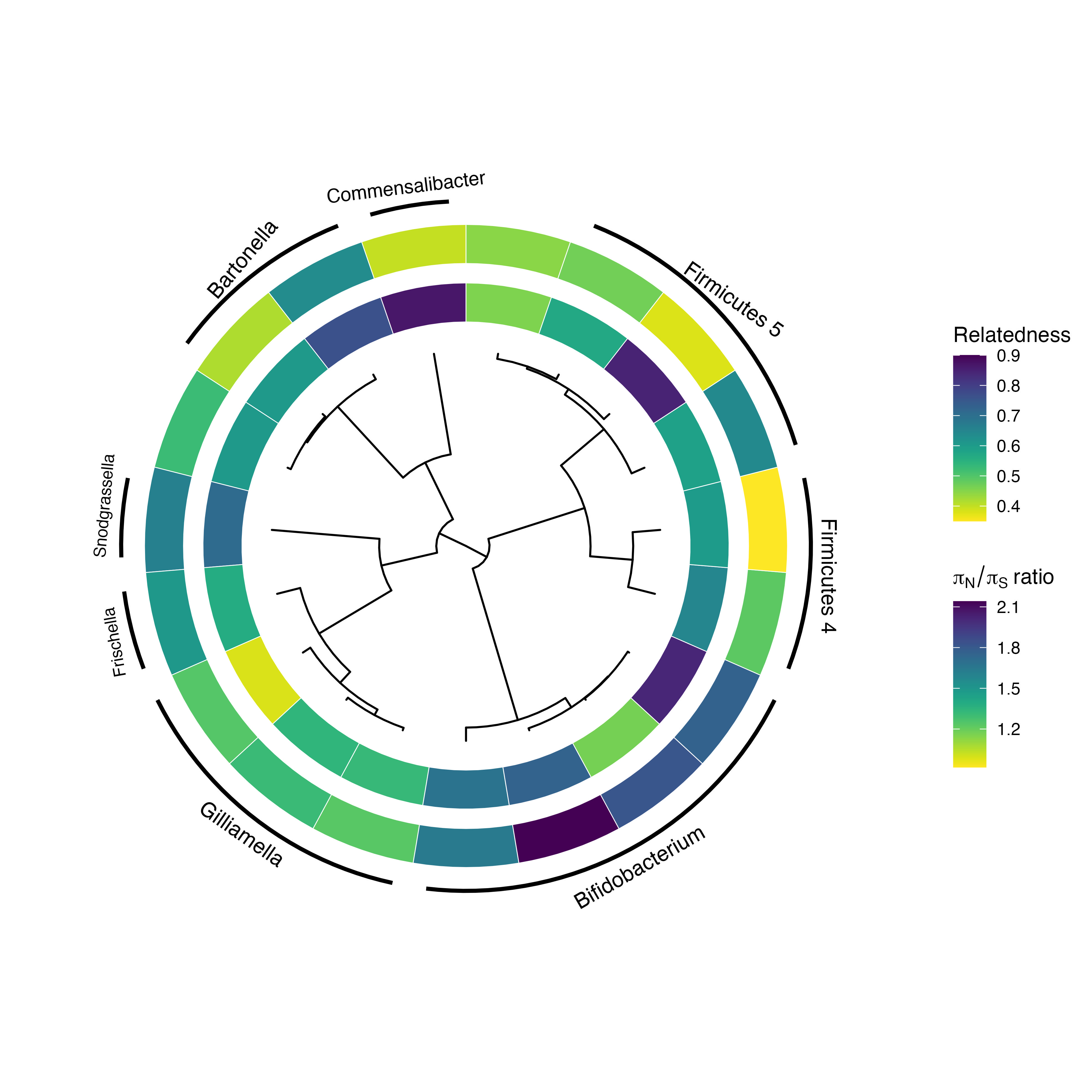

Figure S4: Phylogeny of the 19 species, with metagenomic relatedness (inner ring) and the ratio of non-synonymous to synonymous polymorphism in genes for cooperative traits relative to genes for private traits of the same length (outer ring) shown in colour. Arcs mark the genera.

### Supplement S5: Replication in three more populations

#### Methods

We describe the samples, processing and the calculation of relatedness and polymorphism in the main Methods. Here we describe how we matched MIDAS2 (Zhao et al., 2023) species to our species.

To link MIDAS2 species to our species, we matched each of our reference genomes to a GTDB (Parks et al., 2022) representative genome with skani (v0.4.0; Shaw & Yu, 2023) at an average nucleotide identity of at least 95%. We transferred our labels of genes for cooperative and private traits to each representative genome by best DIAMOND hits (Buchfink, Reuter & Drost, 2021) with at least 90% identity covering at least 80% of both proteins. Genes for private traits took the PSORTb prediction of the reference gene they matched. This was possible for 15 of our 19 species. Barto-2 and Bifido-1.2 had no matching GTDB genome, and for Barto-3 and Firm4-1 the gene identifiers in our SOCfinder output did not match those in the proteomes.

#### Results

Genes for cooperative traits had more non-synonymous relative to synonymous polymorphism than genes for private traits in most species in every population (Figure S5; Table S6). In the three new populations, the ratio was higher in 12 of 14 species in Les Droites 2015, 11 of 13 in Les Droites 2016 and 11 of 15 in Le Grammont 2016, with medians of 1.21, 1.30 and 1.20. Across the three populations, the ratio was 1.17 (95% CI = 0.92–1.49; phylogenetic mixed model with species and population as random effects). With the OH colony reanalysed by the same pipeline, the ratio was higher in 12 of 15 species (median 1.24).

Relatedness ranged from −0.15 to 0.95, with medians of 0.51 to 0.64 per population. The one negative estimate (Gilli-2 in Le Grammont 2016) means that diversity within those bees was slightly higher than in all bees pooled, which can happen by chance when relatedness is close to zero. In the OH colony, relatedness estimated with MIDAS2 agreed closely with our original estimates (Spearman’s ρ = 0.87, n = 15 species).

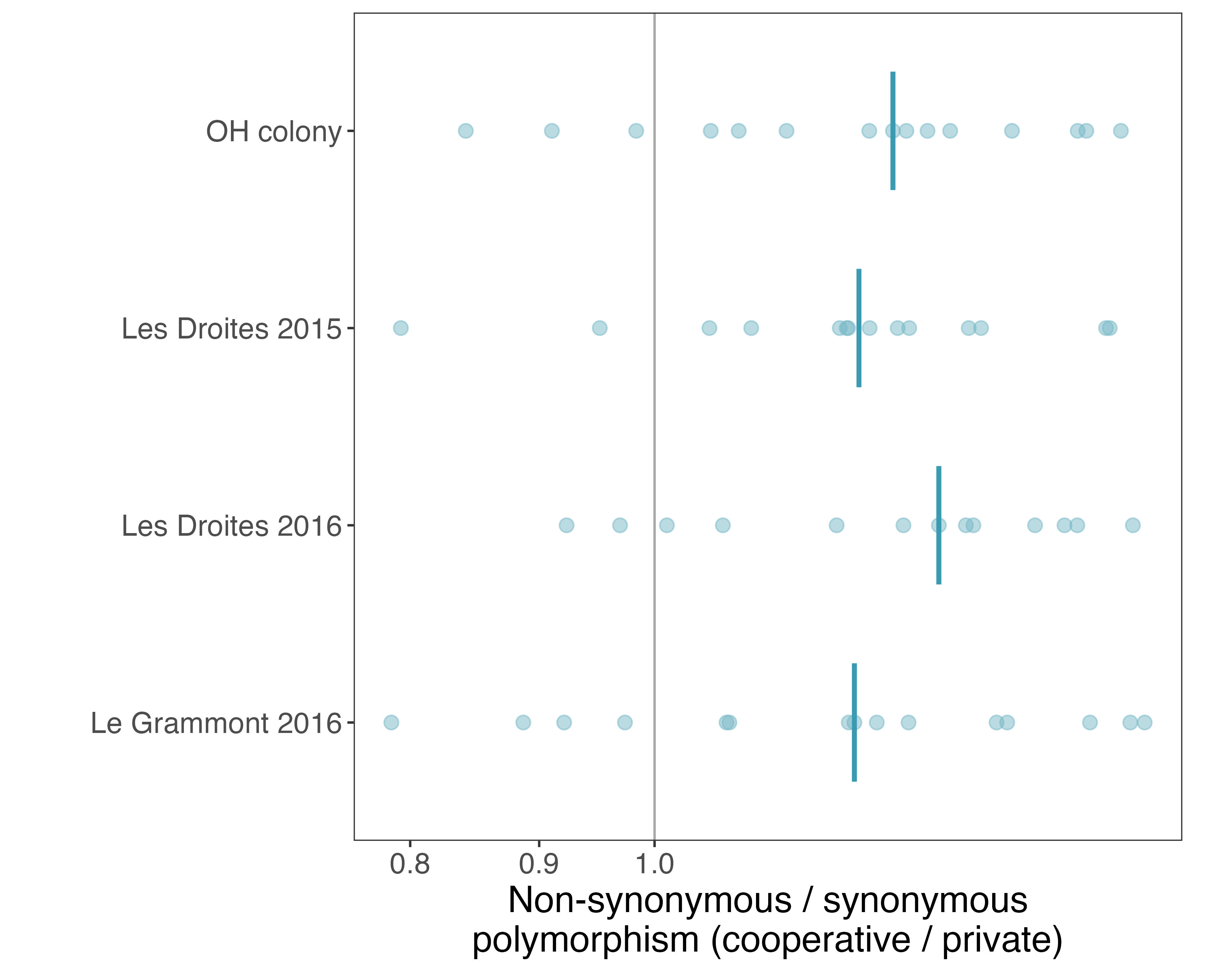

Figure S5: Non-synonymous relative to synonymous polymorphism of genes for cooperative and private traits in four populations of honey bees. Each point is one species in one population (13 to 15 species per population): the ratio of πN/πS in genes for cooperative traits to that in genes for private traits, from a model with gene length as a covariate. The bar shows the median for each population, and values to the right of the grey line at 1 mean that genes for cooperative traits have more non-synonymous relative to synonymous polymorphism. We reanalysed the OH colony with the same pipeline as the other populations, so its values differ slightly from Figure 3.

### Supplement S6: Further checks of the main results

#### Methods

For 1,409 orthogroups, codeml (PAML v4.10.10; Yang, 2007) could not run because the gene tree contained nodes with more than two branches, which arise when strains are nearly identical. We resolved each of these nodes into successive two-way splits with zero-length branches, in arbitrary order, and reran codeml. All 1,409 orthogroups then ran.

We repeated each model of one value per species with the GTDB phylogeny (Supplement S4) in place of our main phylogeny. For dN/dS between strains and positive selection, we also fitted models with every orthogroup as a data point, with the same covariates as in the main models, and phylogeny and species as random effects. For positive selection, which is binary, we fixed the residual variance at 1 and ran these chains for 3.1 × 10⁶ iterations with thinning of 3,000.

#### Results

The results were the same with the GTDB phylogeny and with every orthogroup as a data point (Table S11).

Table S11: Genes for cooperative traits relative to genes for private traits, with three choices of model. Values are ratios (polymorphism, dN/dS) or odds ratios (positive selection, gene loss), with 95% credible intervals and pMCMC. The replication is reported as an effect size only.

| **Signature (species)** | **Main model** | **GTDB phylogeny** | **Every orthogroup** |
| --- | --- | --- | --- |
| πN/πS, OH colony (19) | 1.36 (1.08–1.65), 0.016 | 1.36 (1.11–1.67), 0.013 |  |
| πN/πS, three new populations (15) | 1.17 (0.92–1.49) | 1.17 (0.93–1.51) |  |
| dN/dS between strains (10) | 1.49 (1.14–1.90), 0.008 | 1.49 (1.16–1.88), 0.009 | 1.52 (1.24–1.81), < 0.001 |
| Positive selection (10) | 2.12 (1.29–3.49), 0.011 | 2.10 (1.21–3.23), 0.015 | 2.12 (1.50–2.96), < 0.001 |
| Gene loss (16) | 3.57 (1.41–7.93), 0.015 | 3.58 (1.59–8.26), 0.005 |  |
| dN/dS against the outgroup (13) | 1.51 (0.92–2.48), 0.085 | 1.49 (0.92–2.40), 0.084 |  |

dN/dS between strains was higher in genes for cooperative traits in 9 of the 10 species, and lower in Bifido-1.4. The largest differences were in Gilli-2, Bifido-1.1 and Snod-Amel, where dN/dS in genes for cooperative traits was 1.7 to 2.2 times that in genes for private traits (Table S4). Orthogroups for cooperative traits were more likely than those for private traits to have positively selected codons in all 10 species, with the largest differences in Bifido-1.1, Bifido-1.2 and Bifido-1.4.

Recombination was common, and was detected in 68% of orthogroups for cooperative traits and 55% of those for private traits. We therefore included recombination as a covariate in all models of positive selection.

A higher percentage of orthogroups for cooperative traits than for private traits had been lost in 10 of the 16 species, and a lower percentage in the other six. The difference was largest in Bifido-1.1, Firm5-4, Bifido-1.2 and Bifido-1.4, where 23% to 71% of orthogroups for cooperative traits had been lost, compared with 5% to 49% of those for private traits (Table S3).

In the comparison with the outgroup, synonymous sites were saturated (median dS above 1) in 11 of the 19 species. dN/dS was higher in genes for cooperative traits in 12 of the 13 species with at least five usable genes for cooperative traits. The largest differences were in Bifido-1.1, Bifido-1.2 and Bifido-2, where dN/dS in genes for cooperative traits was 2.2 to 3.6 times that in genes for private traits (Table S5).

### Supplement S7: Functions of lost and positively selected genes

#### Methods

We annotated each reference proteome with eggNOG-mapper (v2.1.13, DIAMOND mode, eggNOG database v5.0.2; Huerta-Cepas et al., 2019; Cantalapiedra et al., 2021). We gave each orthogroup the Gene Ontology (GO) terms of its reference genes, and we used the GO release of 26 July 2026 for term names (The Gene Ontology Consortium, 2025). We tested whether orthogroups that had been lost, or that had positively selected codons, were enriched for particular GO terms, following Belcher & Kelly (2026). We used a hypergeometric test against a background of all orthogroups with at least one GO term, pooled over species, tested terms that annotated 10 to 500 orthogroups, and corrected for multiple tests with the Benjamini–Hochberg method.

#### Results

Orthogroups lost on the strain trees were enriched for carbohydrate metabolism and transport (46 terms at a false discovery rate below 0.05, for example carbohydrate metabolic process, 3.5-fold enrichment, adjusted p < 0.001; Table S8). This enrichment fits the idea that strains of the same species differ in which sugars from the bee’s diet they can use (Ellegaard & Engel, 2019; Li et al., 2026). Only 8 of the 74 lost orthogroups for cooperative traits had GO terms, too few to test separately.

Orthogroups with positively selected codons (FEL) were not enriched for any GO term. The orthogroups for cooperative traits with positively selected codons are listed in Table S9.

### Supplement S8: Metagenomic relatedness and the other signatures of kin selection

Metagenomic relatedness did not predict the difference between genes for cooperative and private traits in any signature, with either phylogeny (Table S12). For non-synonymous relative to synonymous polymorphism across the four populations, we used relatedness in each population, estimated with MIDAS2. Metagenomic relatedness also did not predict the number of genes for cooperative traits, with either phylogeny.

Table S12: Effect of metagenomic relatedness on each signature of kin selection and on the number of genes for cooperative traits (posterior mean slope, 95% credible interval and pMCMC).

| **Response (species)** | **Main phylogeny** | **GTDB phylogeny** |
| --- | --- | --- |
| πN/πS, OH colony (19) | 0.08 (−0.63 to 0.87), 0.84 | 0.08 (−0.69 to 0.82), 0.83 |
| πN/πS, four populations (15) | 0.06 (−0.20 to 0.35), 0.66 | 0.06 (−0.21 to 0.34), 0.66 |
| dN/dS between strains (10) | −0.26 (−1.57 to 1.08), 0.69 | −0.25 (−1.56 to 1.10), 0.70 |
| Positive selection (10) | −0.21 (−2.88 to 2.62), 0.88 | −0.17 (−2.82 to 2.75), 0.90 |
| Gene loss (16) | −0.85 (−3.83 to 2.30), 0.52 | −0.92 (−3.86 to 2.10), 0.50 |
| dN/dS against the outgroup (13) | −0.01 (−1.79 to 1.85), 0.98 | −0.05 (−1.86 to 1.70), 0.94 |
| Number of genes for cooperative traits, SOCfinder (19) | −0.11 (−0.99 to 0.73), 0.80 | −0.13 (−0.98 to 0.67), 0.76 |
| Proportion of genes for cooperative traits, SOCfinder (19) | −0.11 (−0.95 to 0.71), 0.79 | −0.12 (−0.95 to 0.64), 0.74 |
| Number of genes for cooperative traits, Simonet & McNally (19) | 0.55 (−0.54 to 1.58), 0.30 | 0.56 (−0.47 to 1.62), 0.30 |

### Supplement S9: Weapons and carbohydrate-active enzymes

#### Methods

We counted weapons, which are traits that harm or kill competing cells (Granato, Meiller-Legrand & Foster, 2019), and carbohydrate-active enzymes in the reference genome of each of the 19 species, using current databases. The weapons were bacteriocins, which are protein toxins (Sharp & Foster, 2025), and the effectors of type VI secretion systems, which inject toxic proteins into neighbouring cells (Smith et al., 2020).

To find enzymes that break down complex carbohydrates from pollen and nectar, we used run_dbCAN (v5.2.9, database release 5.2.9; Zheng et al., 2023). We searched each proteome with HMMER (v3.4; Eddy, 2011) against the dbCAN profiles and with DIAMOND (v2.2.8; Buchfink, Reuter & Drost, 2021) against CAZy (Drula et al., 2022), and counted a protein only when both searches identified it as a carbohydrate-active enzyme. Because glycoside hydrolases and polysaccharide lyases can act on dietary polysaccharides outside the cell, we analysed only these two classes.

To find bacteriocins, we searched each proteome against the BAGEL4 databases (van Heel et al., 2018) at BAGEL4’s own thresholds. We used BLASTP (v2.17; Camacho et al., 2009) against the class I, II and III bacteriocin databases, with E-value cut-offs of 10⁻⁵, 10⁻⁷ and 10⁻²¹, and hmmsearch against the BAGEL4 bacteriocin profiles with a domain score of at least 25. We searched annotated proteins only, so very short bacteriocin genes that the gene caller missed would not be found.

To find effectors of type VI secretion systems, we searched each proteome with DIAMOND against SecReT6 (v3.0; Zhang et al., 2023). We counted a protein as an effector if it matched a known effector with at least 70% identity over at least 80% of the query protein (E ≤ 10⁻⁵). Counts under a looser threshold (30% identity, 50% coverage) are in Table S2. We identified secretion systems with MacSyFinder (v2.1.6; Néron et al., 2023) and the TXSScan models (Abby et al., 2016).

#### Results

SOCfinder does not include weapons in its list of genes for cooperative traits. Of the 142 weapon genes that had a SOCfinder label, 8 were labelled as being for cooperative traits and 134 for private traits.

We found no effect of metagenomic relatedness on the number of carbohydrate-active enzymes (glycoside hydrolases and polysaccharide lyases; MCMCglmm; posterior mean = 0.005, 95% CI = −1.49 to 1.47, pMCMC = 0.95), or on the number of genes for cooperative traits when we included weapons (posterior mean = −0.07, 95% CI = −0.88 to 0.80, pMCMC = 0.87; Table S2). The results were the same with the GTDB phylogeny (pMCMC = 0.97 and 0.85).

### Supplement S10: Phylogenetic power analysis

To test whether our study could have detected the effect that Simonet & McNally (2021) found, we carried out a phylogenetic power analysis. We asked whether an effect of the same size as in their study of 101 species would be detectable in our study of 19 species. In the data of Simonet & McNally, the relationship between relatedness and the number of genes for cooperative traits has a slope of 28.922 and an intercept of 25.531, and the 95% confidence interval of the slope is 1.191 to 56.65. In each of 1,000 simulations, we drew a slope from a normal distribution with mean 28.922 and standard deviation 14.147, which matches this confidence interval. For each species, we multiplied this slope by the metagenomic relatedness of the species and added the intercept, which gave a simulated number of genes for cooperative traits. We fitted a phylogenetic MCMCglmm model (Hadfield, 2010) to these simulated numbers (Poisson, with the log number of other genes as a covariate), and recorded whether relatedness had a significant effect (pMCMC < 0.05). We used the same phylogeny, prior and chain lengths as in our main models. Relatedness had a significant effect in 57% of the simulations.

### Supplement S11: Relatedness estimated from polymorphism

If selection on genes for cooperative traits is relaxed in proportion to relatedness, their non-synonymous relative to synonymous polymorphism is expected to be 1/r times that of genes for private traits (Figure S1C). We therefore estimated relatedness as 1 divided by the ratio of non-synonymous to synonymous polymorphism in genes for cooperative traits relative to genes for private traits. Across the 19 species in the OH colony, the ratio of 1.36 (95% CI = 1.08–1.65) gives a relatedness of 0.73 (95% CI = 0.61–0.92). In the three further populations, the ratio of 1.17 (95% CI = 0.92–1.49) gives a relatedness of 0.85 (95% CI = 0.67–1.09).

We also estimated relatedness for each species from its own ratio (Table S13). These estimates ranged from 0.47 (Bifido-1.4) to 1.09 (Firm4-2), with a median of 0.77. They are less precise than the estimate across species, and two species (Firm4-2 and Firm5-2) had a ratio below 1, which gives a relatedness above 1. Relatedness from polymorphism was higher than metagenomic relatedness in 14 of 19 species. The two estimates were not correlated across species (Spearman’s ρ = −0.04, p = 0.88), which is another way of showing that metagenomic relatedness did not predict the ratio (Figure 5).

To test whether relatedness from polymorphism predicted the number of genes for cooperative traits, we repeated the three models of the number and proportion of genes for cooperative traits with relatedness from polymorphism in place of metagenomic relatedness, with the two values above 1 set to 1. These models are not circular, because the numbers of genes for cooperative traits are not used to estimate relatedness from polymorphism. We did not test relatedness from polymorphism against the signatures of kin selection, because it is calculated from one of them. Relatedness from polymorphism did not significantly predict the number of genes for cooperative traits found by SOCfinder (MCMCglmm; posterior mean = −0.95, 95% CI = −2.02 to 0.01, pMCMC = 0.07), the proportion of genes that were for cooperative traits (posterior mean = −0.94, 95% CI = −1.89 to −0.002, pMCMC = 0.06), or the number found with the method of Simonet & McNally (posterior mean = −0.49, 95% CI = −1.69 to 0.59, pMCMC = 0.37). The results were similar with the GTDB phylogeny (pMCMC = 0.09, 0.07 and 0.38; Table S7).

Table S13: Relatedness estimated from polymorphism in each species (1 divided by the ratio in Table S10, with 95% confidence interval), and metagenomic relatedness.

| **Species** | **Relatedness from polymorphism (95% CI)** | **Metagenomic relatedness** |
| --- | --- | --- |
| Barto-1 | 0.94 (0.68–1.29) | 0.60 |
| Barto-2 | 0.64 (0.45–0.93) | 0.76 |
| Barto-3 | 0.77 (0.55–1.06) | 0.61 |
| Bifido-1.1 | 0.55 (0.41–0.74) | 0.46 |
| Bifido-1.2 | 0.57 (0.41–0.78) | 0.84 |
| Bifido-1.4 | 0.47 (0.34–0.64) | 0.73 |
| Bifido-2 | 0.61 (0.43–0.86) | 0.69 |
| Commensalibacter | 0.98 (0.71–1.34) | 0.87 |
| Firm4-1 | 0.82 (0.51–1.31) | 0.65 |
| Firm4-2 | 1.09 (0.65–1.83) | 0.60 |
| Firm5-1 | 0.64 (0.42–0.98) | 0.59 |
| Firm5-2 | 1.02 (0.66–1.57) | 0.85 |
| Firm5-3 | 0.85 (0.56–1.29) | 0.57 |
| Firm5-4 | 0.88 (0.59–1.33) | 0.46 |
| Frischella-Amel | 0.67 (0.51–0.89) | 0.56 |
| Gilli-1 | 0.81 (0.63–1.02) | 0.38 |
| Gilli-2 | 0.76 (0.58–1.00) | 0.54 |
| Gilli-3 | 0.81 (0.61–1.08) | 0.53 |
| Snod-Amel | 0.62 (0.49–0.79) | 0.71 |
